# The origin and evolution of mitochondrial tropism in *Midichloria* bacteria

**DOI:** 10.1101/2022.05.16.490919

**Authors:** Anna Maria Floriano, Gherard Batisti Biffignandi, Michele Castelli, Emanuela Olivieri, Emanuela Clementi, Francesco Comandatore, Laura Rinaldi, Maxwell Opara, Olivier Plantard, Ana M. Palomar, Valérie Noël, Amrita Vijay, Nathan Lo, Benjamin L. Makepeace, Olivier Duron, Aaron Jex, Lionel Guy, Davide Sassera

**Affiliations:** Department of Biology and Biotechnology “L. Spallanzani”, University of Pavia, Pavia, Italy; Faculty of Science, University of South Bohemia, České Budějovice, Czech Republic; Department of Clinical, Surgical, Diagnostic and Pediatric Sciences, University of Pavia, Pavia, Italy; Istituto Zooprofilattico Sperimentale della Lombardia e dell’Emilia-Romagna, Pavia Department, Strada Campeggi 59/61, 27100, Pavia, Italy; Department of Biomedical and Clinical Sciences L. Sacco, Pediatric Clinical Research Center “Romeo ed Enrica Invernizzi”, University of Milan, Milan, Italy; Department of Veterinary Medicine and Animal Production, University of Naples Federico II, CREMOPAR Regione Campania, Naples, Italy; Zoonotic Parasites Research Group, Department of Parasitology and Entomology, Faculty of Veterinary Medicine, University of Abuja, Abuja Nigeria; INRAE, Oniris, BIOEPAR, Nantes, France; Center of Rickettsiosis and Arthropod-Borne Diseases (CRETAV), San Pedro University Hospital - Center of Biomedical Research from La Rioja (CIBIR), Logroño, Spain; MIVEGEC (Maladies Infectieuses et Vecteurs: Ecologie, Génétique, Evolution et Contrôle), Univ. Montpellier (UM) - Montpellier, France; Population Health and Immunity Division, The Walter and Eliza Hall Institute of Medical Research, Parkville 3052, Australia; School of Life and Environmental Sciences, University of Sydney, Sydney, NSW 2006, Australia; Institute of Infection, Veterinary & Ecological Sciences, University of Liverpool, Liverpool, United Kingdom; Centre of Research in Ecology and Evolution of Diseases (CREES), Montpellier, France, Montpellier, France; Faculty of Veterinary and Agricultural Sciences, The University of Melbourne, Melbourne, Australia; Department of Medical Biochemistry and Microbiology, Science for Life Laboratories, Uppsala University, Uppsala, Sweden

## Abstract

*Midichloria* are intracellular bacterial symbionts of ticks. Some representatives of this genus have the unique capability to colonize mitochondria in the cells of their hosts. Hypotheses on the nature of this interaction have proven difficult to test, partly due to a lack of data. Indeed, until now, mitochondrial tropism information and genomes were available only for symbionts of three and two tick host species, respectively. Here we analyzed the mitochondrial tropism of three additional *Midichloria* and sequenced nine novel genomes, showing that the tropism is pnon-monophyletic, either due to losses of the trait or multiple parallel acquisitions. Comparative genome analyses support the first hypothesis, as the genomes of non-mitochondrial symbionts appear to be reduced subsets of those capable of colonizing the organelles. We detect genomic signatures of mitochondrial tropism, showing a set of candidate genes characteristic of the strains capable of mitochondrial colonization. These include the type IV secretion system and the flagellum, which could allow the secretion of unique effectors, direct interaction with, or invasion of the mitochondria. Other genes, including putative adhesion molecules, proteins possibly involved in actin polymerization, cell wall and outer membrane proteins, are only present in mitochondrial symbionts. The bacteria could use these to manipulate host structures, including mitochondrial membranes, in order to fuse with the organelles or manipulate the mitochondrial network.

## Introduction

“*Candidatus* Midichloria” (hereafter *Midichloria*, order *Rickettsiales*, class *Alphaproteobacteria*) are bacterial endosymbionts of ticks with the unique capability of mitochondrial colonization. *Midichloria* were first described in the ovaries of the hard tick *Ixodes ricinus* (Lewis 1979; Sacchi et al. 2004), where they are highly prevalent and abundant (Sassera et al. 2006; Sassera et al. 2008), indicating a probable mutualistic interaction. The specific role of *Midichloria* is not known. However, genome sequencing of two *Midichloria* strains (Sassera et al. 2011), and the presence of these bacteria in different host tissues, led to the prediction of multiple roles for the bacteria in detoxification, vitamin provision and energy-related functions (Olivieri et al. 2019). In addition, *Midichloria* symbionts have been detected using molecular techniques, with varying prevalence, in multiple tick species belonging to different genera, with evidence of only partial co-cladogenesis with the hosts (Epis et al. 2008; Montagna et al. 2013; Cafiso et al. 2016; Duron et al. 2017; Buysse and Duron 2018; Al-Khafaji et al. 2019). This pattern suggests a facultative mutualistic behaviour and the capability of these bacteria to be transferred horizontally, in addition to their demonstrated transovarial transmission (Lo et al. 2006; Sassera et al. 2006).

*Midichloria* has been investigated ultrastructurally in three tick species. Although the symbionts in *Ixodes holocyclus* appear to be confined to the cytoplasm (Beninati et al. 2009), in ovarian cells of *I. ricinus*, around 10% of the bacteria occur in the intermembrane space of mitochondria (Sacchi et al. 2004; Comandatore et al. 2021). This intramitochondrial tropism has been reported in *Midichloria* of *Rhipicephalus bursa* (Epis et al. 2008) as well.

The mechanisms and functions of such a unique interaction have been the subject of speculation. Initially, Sacchi and colleagues hypothesized that *Midichloria mitochondrii*, the symbiont of *I. ricinus*, consumed mitochondria with a behaviour reminiscent of predatory bacteria (Sacchi et al. 2004). Comandatore and colleagues found this predatory hypothesis inconsistent with mathematical modelling of quantitative transmission electron microscopy (TEM) data and proposed that *Midichloria* could move within the mitochondrial network instead of preying on the organelles (Comandatore et al. 2021).

Considering that *Midichloria* are found in tick species worldwide and that the mitochondrial tropism is not always present, we reasoned that a comparative, multidisciplinary approach could be used to gather information on the mitochondrial tropism and its evolution. We thus analyzed the intracellular behaviour of additional *Midichloria* strains through TEM and sequenced corresponding genomes to detect genomic signatures of mitochondrial tropism.

## Materials and methods

### Sample collection and preparation

Individuals of the tick species *I. ricinus, Ixodes aulacodi, Ixodes holocyclus, Ixodes frontalis, Hyalomma marginatum* and *Hyalomma scupense* were collected from different geographical regions (Italy, France, Spain, Scotland, Australia, Nigeria) between 2016 and 2019. Ovaries and Malpighian tubules were dissected from freshly collected ticks following a published protocol (Olivieri et al. 2019) for microscopy or genomic analyses, while preserved samples were used whole for genomics only.

DNA was extracted from ovaries or Malpighian tubules (tissues known to harbour the symbionts) of freshly collected female ticks or the whole bodies of larvae, nymphs, and preserved females representing 300 total samples. DNA extraction was performed for each sample using one of the following kits: the NucleoSpin® Tissue Kit (Macherey Nagel, Duren, Germany), the NucleoSpin® Plant II Kit (Macherey Nagel, Duren, Germany), or the Qiagen DNeasy Blood & Tissue Kit (Qiagen, Hilden, Germany). DNA was eluted in sterile water and quantified using an Agilent Tapestation (Santa Clara, California, United States) or Invitrogen Qubit Fluorometer by ThermoFisher (Waltham, Massachusetts, United States). In order to select samples for genome sequencing, the presence and abundance of *Midichloria* were determined by qualitative and quantitative PCR (qPCR). The *Midichloria* of *I. ricinus, I. aulacodi* and *Hyalomma* species were quantified by qPCR assays targeting *cal* and *gyrB (Sassera et al*. 2008, Buysse et al. 2021), while the *Midichloria* of *I. holocyclus* were subjected to qPCR assay targeting the 16S rRNA gene (Beninati et al. 2009).

### Transmission electron microscopy

Dissected ovaries from female ticks (one *Hyalomma marginatum*, two *I. frontalis* and six *Hyalomma scupense*) were embedded in EMbed 812 (epoxy resin) and subjected to TEM observation using a Zeiss EM 900 microscope. Samples were fixed in 0.1 M cacodylate buffer (pH 7.2) containing 2.5% glutaraldehyde for 2 h at 4°C and postfixed in 2% OsO4 in 0.1 M cacodylate buffer for 1.5 h at 4°C. Subsequently, the samples were washed, dehydrated through a progressive ethanol gradient, transferred to propylene oxide, and embedded in Epon 812. Thin sections (80 nm) were stained with saturated uranyl acetate followed by Reynolds’ lead citrate and examined with a Zeiss TEM 900 transmission electron microscope at 80 kV.

### Genome sequencing

Genome sequencing was performed either using short-reads only or a hybrid approach. The *I. holocyclus* sample was sequenced on Illumina MiSeq after Illumina TruSeq Paired-end Library preparation and on a R9.4.1 PromethION flow cell (FLO-PR002) for a 65-hr run using MinKNOW v3.4.6 with default settings. All other samples were sequenced with short reads, either on an Illumina HiSeq2000 using the Illumina HiSeq SBS v3 Reagent Kit for library preparation (*I. frontalis* sample) or on an Illumina HiSeq 2500 after Nextera library preparation (all other samples). FAST5 files containing raw Nanopore signal data were base-called and converted to FASTQ format in real-time using Guppy for PromethION v3.0.5.

DNA from each tick sample was sequenced as a whole extract, thus obtaining mixed reads belonging to the bacteria and the arthropod. Illumina read quality was evaluated with FastQC (Andrews S. 2015), reads were trimmed with Trimmomatic v.0.36 with standard parameters (Bolger et al. 2014) assembled using SPADES v3.10.0 with default parameters (Bankevich et al. 2012) and subjected to the Blobology pipeline (Kumar et al. 2013), thus separating them according to GC content, coverage and taxonomic annotation. Further discarding of contaminating sequences and refining the assemblies was performed manually by analysing the assembly graphs with Bandage (Wick et al. 2015) following a previously published pipeline (Castelli et al. 2019). Long reads were filtered according to BLAST hits against a reference of other *Midichloria* sequences (the published genome of *M. mitochondrii* strain IricVA and the genome of the *Midichloria* symbiont of *H. scupense* sequenced in this work). The short and long reads of *I. holocyclus* were assembled using Unicycler (Wick et al. 2017) with default settings. Genome completeness of all the *Midichloria* genomes was evaluated using miComplete (Hugoson et al. 2020) with the --hmms Bact105 setting.

### Genome annotation

The annotation of the previously published *M. mitochondrii* strain IricVA was manually inspected and revised (see Supplementary file 1 for the manually curated annotation), then used as a reference for gene calling in the preliminary annotation of the novel genomes, using Prokka v. 1.12 (Seemann 2014) with default parameters. Due to the AT-rich nature of endosymbiotic bacterial genomes, automatic annotation of proteins sometimes yields erroneous results, especially for proteins presenting high divergence, such as the secretion system components (Gillespie et al. 2015). The annotations were thus comparatively evaluated and manually standardized using the Prokka annotation and results of BLASTp v.2.2.31 against the nr database. Proteins unique to the intramitochondrial *Midichloria* strains, being absent in the other organisms of the NCBI nt database or showing an amino acid sequence identity lower than 60%, were classified as ‘selected from manual annotation’ for further analyses. We paid special attention to the components of secretion systems, as they are usually involved at the host-symbiont interface (Dale et al. 2002; Gillespie et al. 2015) and can be highly divergent in *Midichloria*. All secretory components not identified by Prokka were searched for by BLASTp using *Escherichia coli* O78 (GCF_020217445.1) sequences as queries. Furthermore, we detected pseudogenes among these components by manually analyzing the sequence alignments of each component and ORFs flanking the annotated component.

### Phylogenetic analyses

A phylogenomic dataset was created using phyloSkeleton (Guy 2017) to include the nine novel *Midichloria* genomes, the previously published *M. mitochondrii* strain IricVA genome, and 35 *Rickettsiales* genomes chosen as representative of the diversity of the order. Orthologous proteins were clustered with OrthoMCL (Fischer et al. 2011) (identity cut-off: 40%, inflation value: 1.1, E value cut-off: 10^−5^), and single-copy panorthologs were aligned with MUSCLE (Edgar 2004). Poorly aligned positions and divergent regions were removed by Gblocks (Castresana 2000). MUSCLE and Gblocks were run with default parameters.

Discordance analysis was performed to remove possible horizontally transferred genes. Phylogenetic trees were inferred for each cluster of single copy panorthologs (RAxML (Stamatakis 2015) using the PROTCATGTR model with 100 rapid bootstraps. The discordance among the phylogenetic trees (Rieseberg 1991; Degnan and Rosenberg 2009) was analyzed with an in-house Perl-based pipeline (Viklund et al. 2012; Guy et al. 2014): all trees were subjected to pairwise comparisons (*i.e*. all possible pairs of phylogenetic trees were compared), resulting in discordance values among pairs of trees; the discordance values were then summed and the clusters of single-copy panorthologs were ranked by decreasing discordance. The discordance between two trees is estimated as the number of highly-supported (bootstrap > 80), incompatible bipartitions divided by the sum of highly supported bipartitions in both trees. A discordance threshold was then set manually to discard orthologs whose origin was putatively a lateral gene transfer event. The final dataset of single copy panorthologs was then concatenated, the most suitable evolutionary model was estimated using modeltest-ng (Darriba et al. 2020) and phylogenetic inferences were repeated in RAxML (Stamatakis 2015) as above.

### Selection of genomic signatures of mitochondrial tropism

In addition to the manual curation of their functional annotation (see above), we applied three methods in parallel to select orthologs potentially involved in mitochondrial tropism.

We applied birth-death models to reconstruct the ancestral state and presence/absence of each orthologs at each node of the *Rickettsiales* phylogenomic tree (maximum likelihood model) using in-house Perl and R scripts and the R package phytools (Revell 2012). We then selected the genes gained and lost in nodes where, based on the phylogeny, the intramitochondrial tropism (IMT) could have originated or been lost (see results).

Intra-mitochondrial symbionts were then compared with non-IMT *Midichloria* and other *Rickettsiales* for gene composition by DAPC (Discriminant Analysis of Principal Components) with the same dataset used in the phylogenomic analysis and ancestral state reconstruction using the R package adegenet (Jombart 2008). The analysis was repeated without the *Midichloria* symbiont of *I. aulacodi*, as no information on the mitochondrial tropism of this strain is available.

A smaller dataset containing a subset of *Midichloria* genomes was used to analyse selective pressures, including just two *I. ricinus* symbiont genomes to increase uniformity and avoid oversampling. A phylogeny for this reduced dataset was inferred as described above (evolutionary model chosen by modeltest: PROTGAMMAJTT). The tree was rooted using the *Rickettsiales* phylogeny previously inferred as a reference and then used for selective pressure analysis of each single-copy ortholog in the dataset. The alignment of each ortholog was analyzed using Hyphy v2.5.8 (Kosakovsky Pond and Frost 2005), considering the aBSREL branch-site model (Smith et al. 2015).

### Prediction of localization of candidate proteins

All protein candidates from the previous analyses (listed in Table S2) were screened with a bioinformatic pipeline to predict their secretion and cellular localization. To detect putative membrane and secreted proteins that could be involved in the interaction between mitochondria and *Midichloria*, we used TMHMM2 (Krogh et al. 2001) for the prediction of transmembrane helices, SignalP5 (Almagro Armenteros et al. 2019) for the detection of proteins secreted through canonical pathways, and SecretomeP (Bendtsen et al. 2005) to search for proteins potentially secreted through non-canonical pathways (all software used with default parameters). Additionally, the candidates were subjected to a screening with tools meant to identify mitochondrial targeting in eukaryotes: DeepLOC in accurate mode (Almagro Armenteros et al. 2017) and TargetP (Emanuelsson et al. 2007). We hypothesized that *Midichloria* could exploit the existing host machinery to enter mitochondria, and hence that the proteins involved in this process would mimic eukaryotic proteins targeted at the mitochondria or have mitochondrial signal peptides (Kunze and Berger 2015).

Finally, to refine predictions and provide hypotheses on their possible roles, all the candidates were manually inspected by accurate re-annotation, BLAST analysis, and NCBI conserved domain search (Lu et al. 2020).

## Results and discussion

### Novel information on mitochondrial tropism

The *Midichloria* endosymbionts of *I. ricinus* and *R. bursa* can localize within host mitochondria (Sassera et al. 2006; Epis et al. 2008), while those of *I. holocyclus* reside only in the cytoplasm (Beninati et al. 2009). We analyzed ovaries of *H. marginatum, I. frontalis* and *H. scupense* with TEM, leading to the observation of two novel intramitochondrial tropism (IMT) strains and one novel non-IMT strain. The symbionts of *I. frontalis* and *H. marginatum* were observed in the cytosol and inside mitochondria (Figure 1A and 1B). Previous work showed that in *H. marginatum* cells, in addition to *Midichloria, Francisella* symbionts can be present at high concentrations (Buysse et al. 2021). The intramitochondrial bacteria herein observed could thus, in theory, be *Francisella*. However, we believe these to be *Midichloria* because i) *Francisella* has never been observed in mitochondria and ii) the observed size and shape are compatible with other *Midichloria* and not with *Francisella (Midichloria*: 0.45×1.2 μm; morphology: bacilli. *Francisella*: 0.2 × 0.2-0.7 μm; morphology: cocci or coccobacilli or rods).

**Figure 1:**
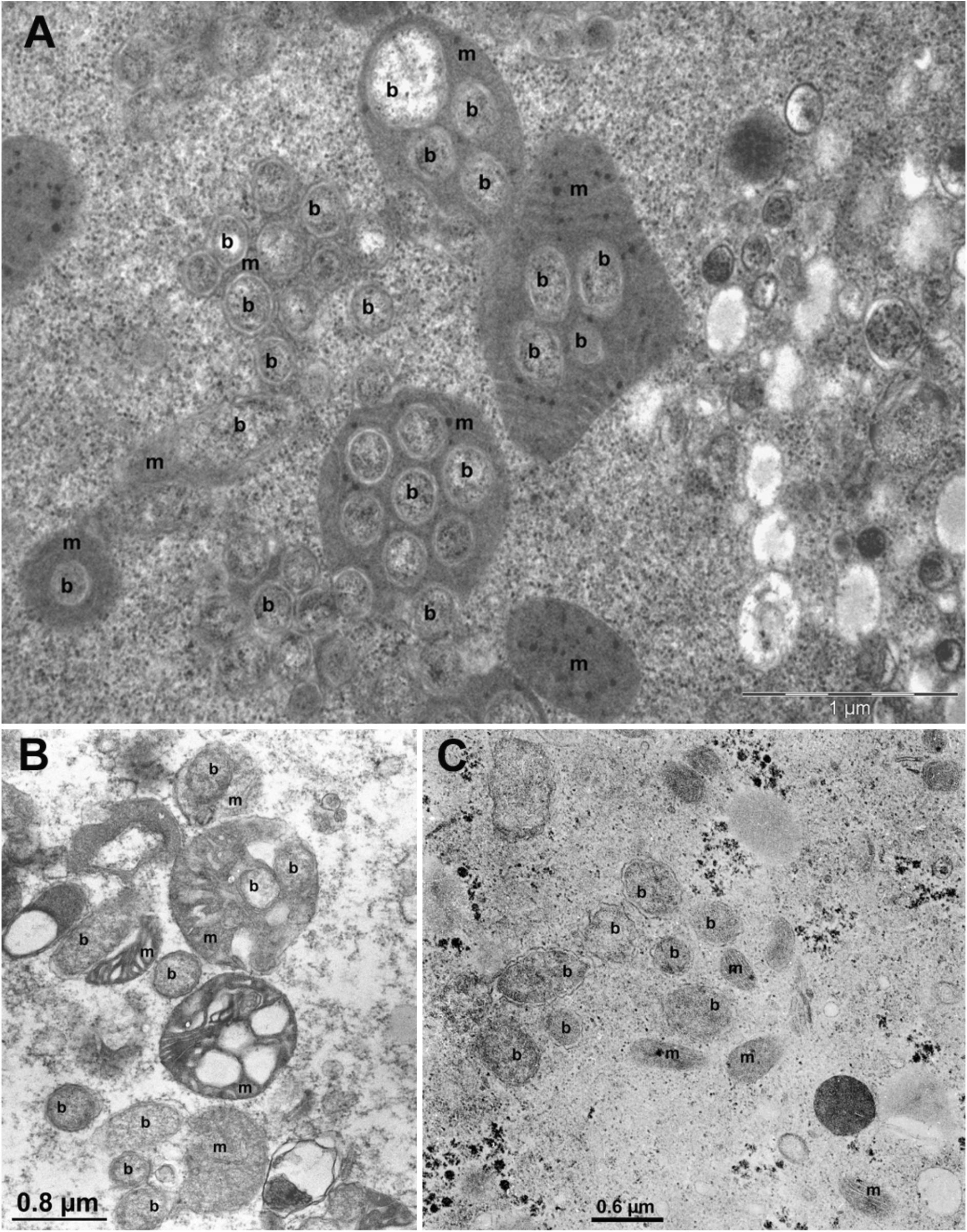
Transmission electron microscopy of oocytes of three tick species showing presence or absence of intramitochondrial tropism of *Midichloria* spp. strains. In figures b indicates bacteria and m mitochondria. A: *Midichloria* sp. symbiont of *Ixodes frontalis* colonizing mitochondria; B: *Midichloria* sp. symbiont of *Hyalomma marginatum* colonizing mitochondria; C: *Midichloria* sp. symbiont of *Hyalomma scupense* in the cytosol, not colonizing mitochondria

The *Midichloria* symbionts of *H. scupense* were observed exclusively in the cytoplasm (Figure 1C) in all six extensively analyzed samples. Lack of evidence of intramitochondrial tropism cannot be conclusively considered evidence of absence. However, quantitative data on *Midichloria* of *I. ricinus* indicate that, on average, around 20% of the total symbionts are intramitochondrial and colonize more than 11% of total organelles (Comandatore et al. 2021). In addition, qualitative data in other species presented here and previously (Epis et al. 2008; Cafiso et al. 2016) show similar levels of “easily detectable intramitochondriality”. We are thus confident in considering the symbiont of *H. scupense* as not capable of colonizing mitochondria. Unfortunately, all efforts to collect ovarian samples of sufficient quality to perform TEM from *I. aulacodi* failed due to the difficulty of sampling and storing high quality samples in the area of distribution of this tick species (*i.e*. West and Central Africa).

### Phylogeny of *Midichloria*

We sequenced nine novel *Midichloria* genomes, estimated based on gene presence to be complete (see Supplementary Table S1 for details) and used them for gene-content and phylogenetic analyses. The topology of the phylogenomics-based tree of the *Rickettsiales* (Figure 2, see also Supplementary Figure 1 for a complete tree) appears consistent with previously published data (Castelli et al. 2016) and shows generally highly support. One exception is a single node within the *Midichloriaceae* family (49% bootstrap support). This is not surprising considering that previous studies showed that the inner topology of *Midichloriaceae* is poorly resolved (Castelli et al. 2016; Giannotti et al. 2022). Nevertheless, this does not affect the robustness of the topology within *Midichloria* (all nodes supported with 100% bootstraps, except for the relationships between the highly similar symbionts of *I. ricinus*), nor the inference on the “phylogenetic surrounding” of *Midichloria*, including the sister-group relation with *Cyrtobacter*.

**Figure 2:**
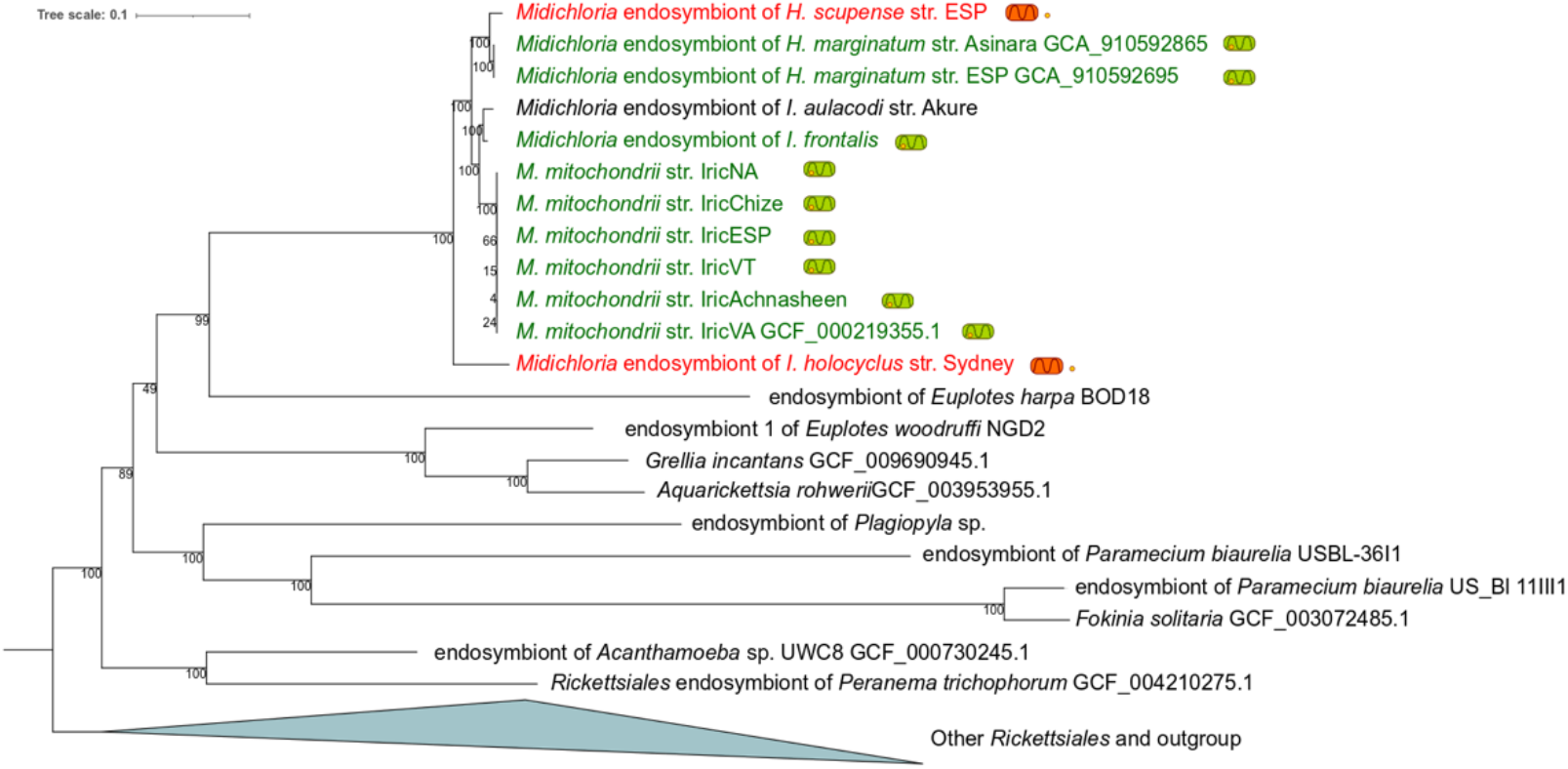
Phylogenomic tree of *Rickettsiales* bacteria, obtained with RAxML, focusing on *Midichloriaceae*. The intramitochondrial *Midichloria* strains are indicated by green mitochondria harbouring a bacterium, while the non-IMT *Midichloria* strains are indicated by a bacterium flanked by orange mitochondria

We find no evidence of full co-cladogenesis between hosts and symbionts, as *Midichloria* symbionts of *Hyalomma* and *Ixodes* ticks do not cluster separately. Considering the monophyly of the symbionts of *Hyalomma*, the previously postulated recent acquisition of *Midichloria* by this clade (Buysse et al. 2021), and that the phylogeny of the symbionts of *Ixodes* ticks appears to be congruent with that of the hosts (D’Amico et al. 2018), the more likely explanation of the presented topology is horizontal transmission from an *Ixodes* species to *Hyalomma*, consistent with previous results (Buysse et al. 2021). The most parsimonious scenario is that of a single transfer event in the ancestor of the *H. scupense* - *H. marginatum* clade, but the possibility that two separate events occurred cannot be excluded.

Another interesting result of the phylogenetic analysis is the sister group position of the *Midichloria* symbiont of the Australian tick *I. holocyclus* to all other *Midichloria*, with a relatively long branch. This Australian tick species has diverged early from the other *Ixodes* and is believed to have been isolated in Australia after the separation from Gondwanan landmasses (99 Myr, Dietrich et al. 2014). Hence, the observed topology suggests that the *Midichloria* symbiosis with *Ixodes* spp. could predate the separation between the radiation of this genus and the separation of Australian and Gondwanan landmasses.

Surprisingly, intramitochondrial tropism is also a non-monophyletic trait. Given the tree topology and the position of the non-IMT *Midichloria* strains, three evolutionary hypotheses for the origin of the intramitochondrial tropism can be drawn: (i) intramitochondrial tropism originated after the early divergence of *Midichloria* of *I. holocyclus* and was lost by the symbiont of *H. scupense*; (ii) intramitochondrial tropism originated before the separation of the symbiont of *I. holocyclus* and the other *Midichloria* and was then lost by the symbionts of *I. holocyclus* and *H. scupense*; (iii) the intramitochondrial tropism evolved independently in the symbionts of *I. ricinus, I. frontalis* and *H. marginatum*.

Hypotheses (ii) and (iii) imply three evolutionary events each, whereas hypothesis (i) implies two and thus is the most parsimonious if gain and loss events are considered equally likely. It must also be considered, however, that many examples, and what is believed to be the general trend of genome evolution in endosymbionts, clearly indicate that gene loss is a relatively common event in these lineages, while the acquisition of a trait is much rarer (Moran et al. 2008). If we consider the event of acquisition of a novel feature much less likely than loss, the discriminant factor would be the number of acquisitions. Based on this reasoning, we considered both hypotheses (i) and (ii) to be similarly likely for the purpose of analyzing gene gains and loss on the phylogenetic tree.

### Genomic signatures of mitochondrial tropism

The nine novel draft genomes of *Midichloria* from seven tick species have sizes ranging from ~0.9 Mb to ~1.2 Mb (Table S1). Non-IMT strains present smaller genomes: 933,600 and 1,064,904 bp for the *I. holocyclus* and *H. scupense* the symbionts, respectively.

To detect potential signatures of mitochondrial tropism, we performed multiple *ad-hoc* analyses to determine differences in the patterns of gene presence and selective pressure between intra-mitochondrial and non-intra-mitochondrial organisms (Supplementary Table S2). We used multiple independent methods to detect possible genes involved in mitochondrial tropism to minimize false negatives, allowing the selection of a higher number of potential candidates. Accordingly, regardless of whether genes were detected by one or multiple methods, we included them in the following analyses.

We manually curated annotations and orthology results to determine the differential presence of orthologs in the genomes. This identified 45 orthologs unique to the IMT *Midichloria* strains or diverging in amino acid sequence (cut-off of identity: 60%) between IMT and non-IMT *Midichloria* or other organisms in the NCBI database. Additionally, discriminant analysis of principal components (DAPC) detected 38 genes characteristic of the IMT *Midichloria* genomes (Supplementary Figure 2). In parallel, we performed an ancestral state reconstruction through birth-death models to reconstruct the gene content of each node of the phylogeny, focusing on the nodes where intramitochondrial tropism could have originated. The analysis indicated that 155 orthologs were gained and 7 lost in the node encompassing all the sequenced *Midichloria* strains (Last *Midichloria* Common Ancestor), and 88 orthologs were gained and 1 lost in the node after the divergence of the *Midichloria* symbiont of *I. holocyclus* from the rest of the genus.

Positive selection was inferred using the adaptive branch-site random effects (aBSREL) model. Only single-copy orthologs were considered in the analysis (560 in total), since the presence of multiple gene copies can affect the results by reducing the signal of negative selective pressures (Roux et al. 2014; Cicconardi et al. 2017). We observed a signal of positive diversifying selection in at least one branch of the *Midichloria* tree for 16 genes.

While no microscopy data are available for the *Midichloria* symbiont of *I. aulacodi*, its genome shares many characteristics with intramitochondrial strains. Its genome size is 1.15 Mb, its gene content is similar to that of IMT strains, and the initial DAPC showed this strain to cluster with IMT strains (Supplementary Figure 2), suggesting it could be IMT as well.

### Proteins potentially involved in intramitochondrial tropism

The genes belonging to the orthogroups selected in the comparative genomic analyses (292 candidate proteins in the reference IricVA strain) were subjected to a bioinformatic pipeline to predict which could be membrane-bound or secreted and which had a signal peptide for mitochondrial targeting, under the hypothesis that these characteristics could be indicative of interaction with mitochondria. Of the 292 proteins subjected to this pipeline, 19 and 59 were predicted to be membrane-bound (TMHMM2) or secreted (23 indicated by SignalP5, 30 by SecretomeP, and 6 by both tools), respectively. In order to obtain additional clues on the possibility of an interaction with the mitochondria, we used two tools designed for the analysis of eukaryotic proteins (DeepLoc and TargetP), as none are available for prokaryotes due to the uniqueness of this tropism. A potential mitochondrial localization was detected in 24 total proteins (23 predicted by DeepLoc, 1 predicted by both tools). As one can expect from an investigation on such a unique tropism, many of the proteins identified through our investigation have unknown functions, while the annotation of the others allows some speculation.

Fourteen candidate proteins containing VCBS or FG-GAP repeats were detected and further investigated. No homologs of these proteins are found in any other *Rickettsiales* outside the genus *Midichloria*, and these gene families are more represented in the intramitochondrial members of the genus. Seven of them present a secretion signal, of which three are also predicted to be localized to mitochondria. These two types of repeats are found in adhesion proteins, even in symbiotic bacteria (Christensen et al. 2020) and FG-GAP repeats are considered to be homologs of integrins, eukaryotic receptors involved in cell adhesion to other cells and matrices (Chouhan et al. 2011). Notably, a *Trypanosoma brucei* protein containing both FG-GAP and VCBS repeats was localized in mitochondria (Namyanja et al. 2019), suggesting that these repeats could be associated with mitochondrial interaction.

Another interesting protein potentially involved in intracellular motility and interaction with mitochondria is septation protein A. This protein is present in all IMT *Midichloria* and absent or pseudogenized in all non-mitochondrial ones. SepA is necessary for intracellular division and spread in *Shigella*, as it allows the bacterium to polymerize host actin (Mac Síomóin et al. 1996). Interestingly, *Rickettsia* has also been reported to be capable of polymerising actin in host cells, although with different molecular mechanisms (Reed et al. 2014) that are absent in *Midichloria*.

The function of fifteen candidate proteins in our dataset, absent in at least one of the two non-intramitochondrial *Midichloria*, appear to be related to the cell wall and outer membrane, including structural components (*e.g*. porins), biosynthetic enzymes (*e.g*. glycosyltransferases putatively involved in lipopolysaccharide synthesis, UTP-glucose-1-phosphate uridylyltransferase), and processing/regulation factors (flippase, AsmA family protein, D-alanyl-D-alanine carboxypeptidase). The loss of these genes in non-IMT *Midichloria* is indicative of an overall reduction in complexity of the external cell layers. This reduction is similarly found in other lineages of intracellular bacteria, including *Rickettsiales*, and is considered one of the reductive steps towards specialization in the interaction with the host (Floriano et al. 2018; Benerjee and Kulkarni 2021; Atwal et al. 2021; Nardi et al. 2021). Interestingly, lipopolysaccharide is also implicated in recognition of the host nucleus by *Holospora*, a bacterium displaying a peculiar intranuclear localization inside host cells (Fujishima 2009). This observation suggests an intriguing potential analogy in the case of *Midichloria* and mitochondria. Moreover, five putative disulphide dehydrogenases were absent in at least one of the two non-IMT *Midichloria*. In contrast to their intramitochondrial relatives, these enzymes may not be required in exclusively cytoplasmic *Midichloria* that do not encounter the levels of oxidative stress present in the intermembrane space of the mitochondria.

The proteins detected by our analyses also include the subunits of the cytochrome C oxidase (*cbb3*) and other genes involved in the synthesis and processing of its prosthetic heme,absent in non-IMT *Midichloria*. Considering the higher affinity for oxygen but lower efficiency of *cbb3* oxidases compared to typical CcO oxidases, the former are more suitable for the colonization of microaerobic environments (Pitcher and Watmough 2004). The presence of *cbb3* in IMT *Midichloria* could be due to its higher affinity for oxygen being an advantage in a hypothetical competition with host mitochondrial oxidases. The absence of *cbb3* in non-IMT *Midichloria* suggests a streamlined oxidative phosphorylation electron transport chain, re-oxidizing the entire quinone pools directly reducing oxygen, with a less efficient cytochrome bd type oxidase (Borisov et al. 2011). Such a pattern appears analogous to what observed in other intracellular bacteria including another member of *Midichloriaceae* (Floriano et al. 2018). Indeed, in addition to providing clues of the means of interaction between IMT symbionts and mitochondria, the candidates described here appear to paint a general picture of streamlining in non-IMT *Midichloria*. Accordingly, these lineages could be experiencing a pattern of genomic and functional reduction towards more classical nutritional symbiosis. Finally, several identified proteins were deemed components of secretion systems or putative effector molecules and are treated in detail below.

### Type 4 secretion system and flagellum are absent in non-IMT *Midichloria*

Based on the results of our analysis, we carefully analyzed the pattern of presence of genes coding for the flagellum and Type 4 secretion systems (T4SS). T4SS machinery is considered one of the characteristic traits of *Rickettsiales* (including *M. mitochondrii*), is vertically inherited from the ancestor of the order, and is likely involved in the delivery of effectors for the interaction with the host cells (Degnan et al. 2010; Gillespie et al. 2010, Schön et al 2021, Rice et al. 2017). We found 12 main components of the typical *Rickettsiales* T4SS machinery, including known paralogs, in IMT *Midichloria* (the only exception being a *VirB2* paralog missing in the symbiont of *I. frontalis*). On the other hand, in the non-IMT symbionts of *H. scupense* and *I. holocyclus*, most components are missing or pseudogenized (Figure 3). Thus, non-IMT *Midichloria* are predicted to lack a functional T4SS, quite a peculiar absence among *Rickettsiales*.

**Figure 3:**
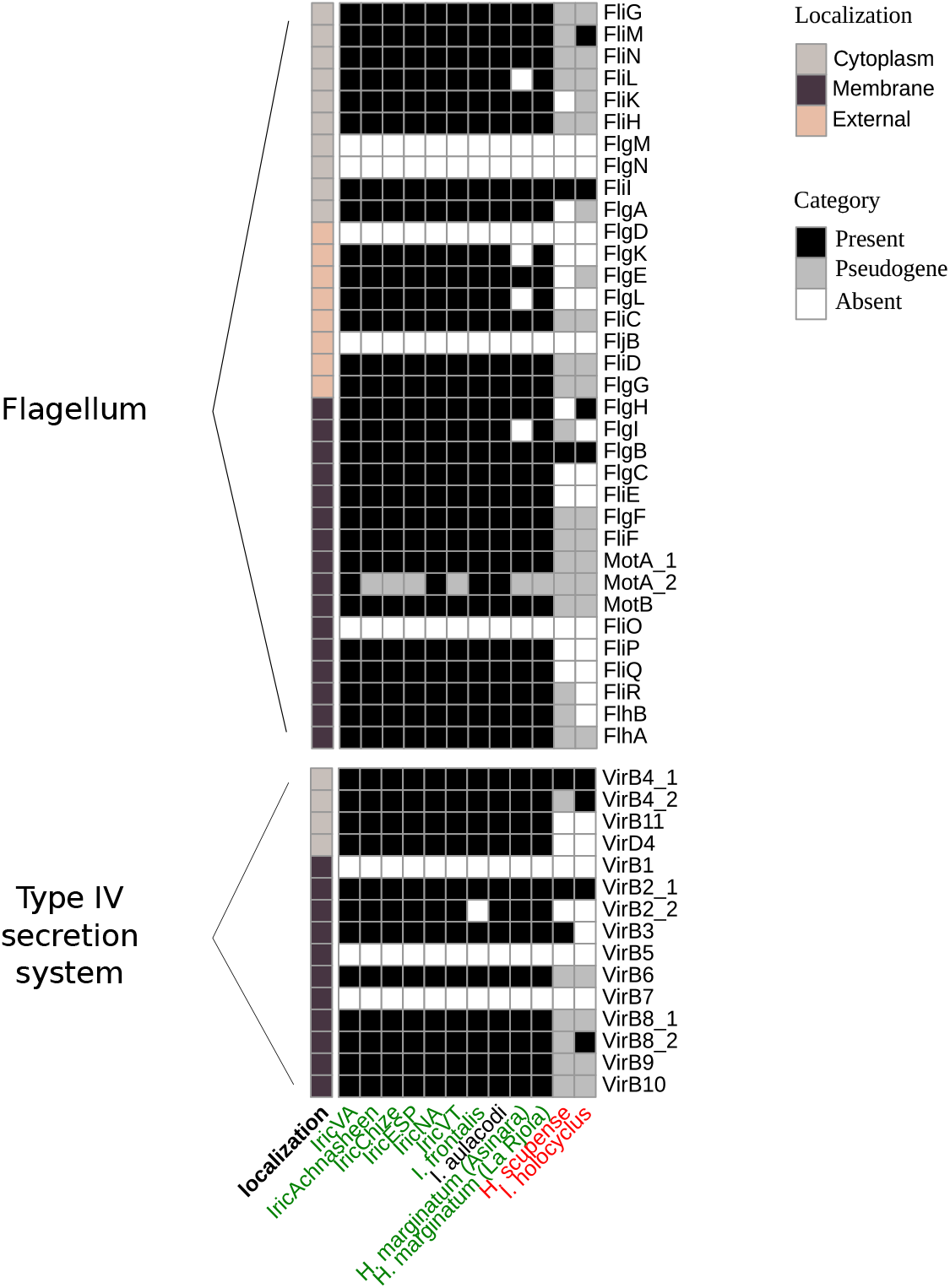
Presence of intact and pseugenized genes encoding for components of the flagellum and of the IV secretion system in the *Midichloria* genomes. Iintramitochondrial symbionts are indicated in green, non-intramitochondrial symbionts in red, the symbiont of *Ixodes aulacodi*, with unknown tropism, is in black.

Among *Rickettsiales* the flagellum is not as common as the T4SS, but it was present in the common ancestor of the order and then independently lost in most lineages (Sassera et al. 2011; Schön et al. 2021). Interestingly, while canonically responsible for movement, this machinery has also been hypothesized to be involved in active invasion of host cells by bacteria (Horstmann et al. 2020), and in the establishment of symbiotic interactions (Shimoyama et al. 2009). Additionally, the transmembrane portion of the flagellum is a Type 3 secretion system (T3SS) homologous to the one found in the injectisome (Cornelis 2006; Diepold and Armitage 2015), the latter being a complex nanomachine involved in the interaction between bacteria and eukaryotic cells (Dale et al. 2002; Cornelis 2006; Abby and Rocha 2012).

When flagellar genes were first identified in *M. mitochondrii*, it was suggested that this macromolecular complex could be involved in horizontal transfer of the symbiont or in interaction with mitochondria (Sassera et al. 2011), a hypothesis corroborated by the fact that no flagella have been observed in *Midichloria* in tick cells (Mariconti et al. 2012). Here we found almost complete gene sets for flagella in all IMT *Midichloria* (Figure 3). Conversely, the two non-IMT genomes display only four of these proteins each (two in common), and thus can be predicted to lack a functional flagellum.

*Midichloria mitochondrii* has an unusually long (“giant”) flagellin (FliC) protein (1039 amino acids). The functional significance of this condition, which evolved convergently in other unrelated bacteria, is still unclear (Thomson et al. 2018). We found that all IMT *Midichloria* have comparably “giant” flagellins, while this protein is absent in non-IMT members of this genus. Additionally, we found two divergent copies of the gene coding for the flagellar stator protein MotA in a non-monophyletic subgroup of four IMT *Midichloria* (two symbionts of *I. ricinus* and those of *I. frontalis* and *I. aulacodi*). In the other six IMT *Midichloria*, also not monophyletic, one of the two copies (which we termed *motA_2*) appears split into two distinct ORFs. Non-IMT *Midichloria* lack *motA* or retain very short and clearly non-functional gene remnants. The uniqueness of these two components and, in general, the divergence of all flagellar components of *Midichloria* from the orthologs in any characterized flagellum bring additional support to the hypothesis that the flagellar machinery in *Midichloria* could have a unique structure and function.

The sharp observed patterns of presence/absence of the flagellum and T4SS within the genus *Midichloria* suggest that these systems may play a role in intramitochondriality. This appears reasonable also considering that T3SS and T4SS are highly important structures at the symbiont-host interface in other bacteria, delivering many effector molecules, such as DNA or proteins, into eukaryotic host cells (Degnan et al. 2010; Gillespie et al. 2010). Furthermore, considering that such systems are vertically inherited from the ancestor of *Rickettsiales* (Gillespie et al. 2010; Sassera et al. 2011b; Schön et al. 2021) and are still present in many members of the order outside the *Midichloria* genus, it seems more plausible to consider them as potential “permissive” traits for the evolution of intramitochondrial tropism, rather than direct determinants *per se*. Thus, alternative and non-mutually exclusive hypotheses can be drawn on how pre-existing flagella and secretion systems could have been involved in the development of IMT. Especially in the case of flagellum/T3SS, it is possible that such a system, initially responsible for motility, acquired new components/functions, considering the presence of additional MotA subunit(s), the giant-sized flagellin or even additional not yet identified components or effectors. Interestingly, studies on *Bartonella* also suggest that the T4SS and T3SS, working in concert with other protein complexes, allow the bacterium adherence and entry into erythrocytes (Eicher and Dehio 2012; Guy et al. 2013).

Focusing on secretion activities, we hypothesize that new dedicated secreted effectors were acquired or co-opted (by gene birth, neofunctionalization or horizontal transfer) for roles relevant to intramitochondriality, such as entry into or survival within mitochondria. Thus, it is interesting to pinpoint which proteins in our dataset could represent potential effectors. In particular, eight of them, absent/pseudogenized in both non-IMTs or at least in the symbiont of *I. holocyclus*, were deemed potential candidates after accurate manual inspection. Six of these proteins contain domains of protein-protein interaction common in host-associated bacteria (Schulz et al. 2016), namely ankyrin-repeats (four proteins), and tetratricopeptide repeats (two proteins). One containing ankyrin repeats (MIDI_RS04925) seems distinctive for having no blast homology outside genus *Midichloria*, while another has several hits in eukaryotes (MIDI_RS03795). Another one of these eight putative effectors (MIDI_RS01890) displays a partial GDP-fucose O-fucosyltransferase domain and an uncharacterized HtrL/YibB-like domain, and its homology (in particular within the HtrL/YibB-like domain) with eukaryotic sequences suggests that, similar to the cases described above, it could be another possible effector mimicking host proteins. Indeed, *Bacterioides fragilis* uses fucosylation to efficiently colonize the human gut (Coyne et al. 2005). Interestingly, the eighth protein (MIDI_RS06020) is homologous to LepB of *Legionella pneumophila*, which is a Rab GTPase-activating protein (GAP) effector important for intracellular vesicular trafficking, and secreted by T4SS (Mihai Gazdag et al. 2013; Mishra et al. 2013; Dong et al. 2016).

Finally, there is no evidence on the timing of the loss of these systems with respect to the supposed loss(es) of IMT among *Midichloria*. It is possible that, similarly to other *Rickettsiales*, such systems in IMT *Midichloria* exert or exterted other functions not related to IMT (*e.g*. delivering different effectors). One example could be in the invasion of new host cells during horizontal transfer, a capability for which there is strong indirect evidence among *Midichloria* (Di Lecce et al. 2018; Cafiso et al. 2019). Indeed in another genus of tick-associated bacteria, the pathogen *Coxiella burnetii* (capable of horizontal transfer) has a larger genome and is equipped with a T4SS. In contrast, mutualistic, endosymbiotic *Coxiella* have smaller genomes and have lost the T4SS (Nardi et al. 2021). Along a similar line of thought, one might question whether a true flagellar motility could be still involved in similar processes, *e.g*. during intermediate passage in a vertebrate (Sassera et al. 2011). It is thus intriguing to hypothesize that non-IMT *Midichloria* may be evolving on a convergent path (*i.e*. strictly vertically transmitted mutualists), leaving open the question of whether intramitochondriality is linked to capability of horizontal transfer.

## Conclusions

Our comparative analyses detected genes potentially involved in the interaction of *Midichloria* with mitochondria. The secretion systems missing in non-IMT symbionts appear highly promising, leading to the hypothesis that IMT *Midichloria* could could exploit these systems for delivering novel effectors and/or could have an alternate, uncharacterized modification of flagellum, possibly involved in mitochondria invasion. Multiple points, in addition to the differential presence, support this last hypothesis: i) flagella have never been seen in *Midichloria* bacteria, ii) flagellar genes of *Midichloria* are highly divergent, iii) the exaptation of flagella into specialized structures associated with host cell invasion has been reported in other bacteria.

The presented results represent a step towards understanding the origin and characteristics of the intramitochondrial tropism of *Midichloria* in ticks, but clearly this work only provides the basis, and requires experimental validation. Nonetheless, a better understanding of the mechanisms of interaction between *Midichloria* and mitochondria could in the future provide a tool to investigate mitochondrial processes and diseases.

Finally, the functional significance of intramitochondriality for *Midichloria* and the host remains an open question that is also critical for understanding the evolutionary processes leading to its origin(s) and its loss(es) during *Midichloria* evolution. Considering that smaller genomes and simplified functionalities of non-IMT *Midichloria* are reminiscent of reduction patterns of nutritional symbionts in ticks and other arthropods, IMT could be seen as a dispensable trait, lost as a side-effect of genomic and functional streamlining, or potentially due to a yet undetermined detrimental effect. From a different perspective, loss of intramitochondriality might also result from an evolutionary cul-de-sac, as a consequence of Muller’s ratchet, in hosts offering unsuitable conditions. Future analyses on datasets, including additional host species and populations, may help to clarify this fascinating point.

## DATA AVAILABILITY

Genomes are available on NCBI under project PRJNA815963.

## ACKNOWLEDGEMENTS

This research was financially supported by the Human Frontier Science Program (HFSP), Young Investigator Program grant RGY-0075 to DS and AJ; by the Italian Ministry of Education, University and Research (MIUR): Dipartimenti di Eccellenza Programme (2018– 2022) Department of Biology and Biotechnology ‘L. Spallanzani’ University of Pavia to DS; by the European Molecular Biology Organization fellowship EMBO-STF-8120 to AMF; AJ is also supported from the Australian National Health and Medical Research Council (APP1194330), Victorian State Government Operational Infrastructure Support and Australian Government National Health and Medical Research Council Independent Research Institute Infrastructure Support Scheme.

## SUPPLEMENTARY MATERIAL

Supplementary Figure 1: Complete phylogenomic tree of *Rickettsiales* bacteria, obtained with RAxML. The intramitochondrial *Midichloria* strains are indicated by green mitochondria harbouring a bacterium, while the non-IMT *Midichloria* strains are indicated by orange mitochondria flanked by a bacterium

Supplementary Figure 2: Principal Component Analysis showing that IMT *Midichloria* (in blue) form a distinct, tight cluster, together with the *Midichloria* symbiont of *I. aulacodi*, for which the tropism could not be assessed (in grey). The rest of the *Rickettsiales* dataset (in red) forms multiple clusters well separated from the IMT cluster, with the two non-IMT *Midichloria* being the points placed closest to the IMT cluster

Supplementary file 1: sequence and manual annotation of the proteins of the *Midichloria mitochondrii* strain IricVA

Supplementary Table 1: datasets with information regarding the analyzed *Midichloria* strains, their tropism (IMT stands for intraMitochondrial) and the characteristics of the genomes

Supplementary Table 2: List of the candidate proteins, including identifiers, annotation and results of the screening processes

## Notes

### Competing Interest Statement

The authors have declared no competing interest.

