## Supplementary figures and images for "The origin and evolution of mitochondrial tropism in *Midichloria* bacteria"

### supplementary Figure 1

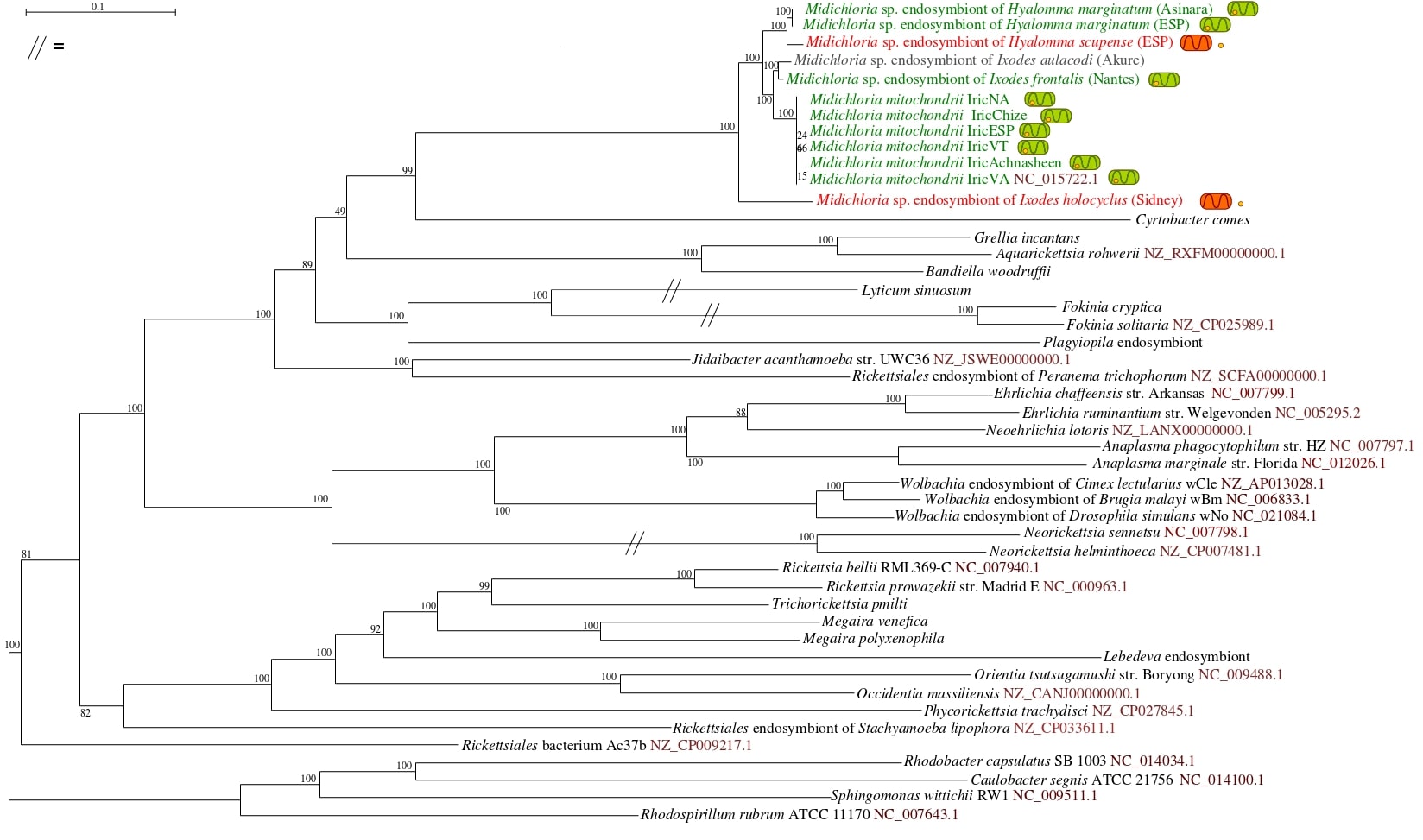

### supplementary Figure 2

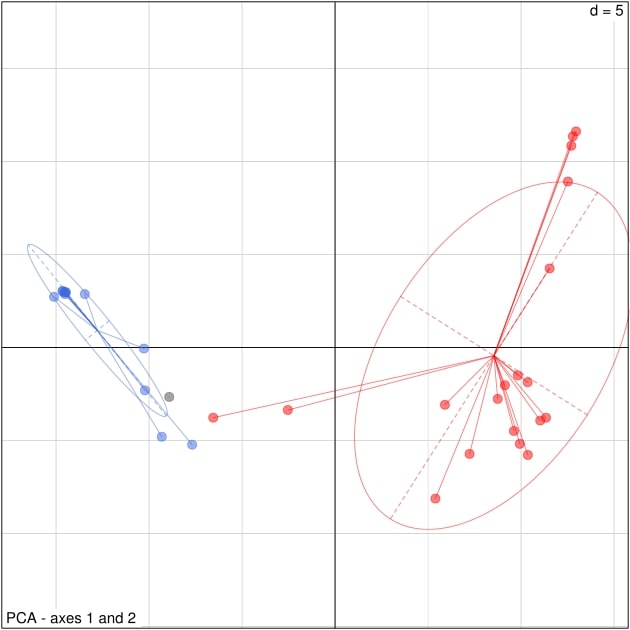
